# Use of anthropogenic material for extended ornamented phenotype in two fairy-wrens

**DOI:** 10.1101/2025.10.06.680756

**Authors:** Jaden Salett, James A. Kennerley, Ryan Jack, Jon Coleman, Michael S. Webster, William E. Feeney, Jordan Boersma

## Abstract

Birds often advertise their quality to potential mates through sexual displays that complement their colorful plumage. Some species use materials to enhance their attractiveness, such as the use of colorful fruits and anthropogenic materials in the display bowers across bowerbirds. Here we report the first observations of anthropogenic materials in sexual displays by two fairy-wren species. On two separate occasions and sites, we witnessed a White-winged and Red-backed male carrying a piece of plastic as a substitute for a flower petal during a petal display, which are used across *Malurus* to enhance reproductive fitness. Given increasing plastic pollution globally, the use of anthropogenic materials as part of extended phenotypes in birds and other animals will likely increase and it will be important to understand the effect this has on populations.

## Introduction

Sexually selected ornaments are well documented across avian taxa, typically involving colourful plumage that aids in attraction of mates (Moller and Pomiankowski 1993). Some passerines further advertise their quality through alterations of the physical environment, i.e. through their extended phenotype (Dawkins 1982; Turner 2004; Schaedelin and Taborsky 2009; Blamires 2013; Wells 2015). For instance, Black Wheatears (*Oenanthe leucura*) signal mate quality by carrying heavy stones to nest sites (Moreno *et al*. 1994), bowerbirds (Ptilonorhynchidae) build and decorate elaborate display arenas (Borgia 1985), and fairy-wrens (Maluridae) present colourful natural materials as nuptial gifts to females (Rowley 1991; Karubian and Alvarado 2003; Boersma *et al*. 2023).

Fairy-wrens (*Malurus* sp.) are model organisms for sexual selection studies owing to their elaborate plumage, sexual display behaviour, and pronounced sexual promiscuity (Dunn and Cockburn 1999; Webster et al., 2007, 2008). Male fairy-wrens will often embark on extra-territorial forays to display to extra-pair females to enhance their reproductive fitness (Rowley 1991; Karubian and Alvarado 2003). Several *Malurus* species often use extended phenotypes, presenting flower petals, berries, or leaves to extra-pair females that complement or contrast with their sexually-selected plumage (Rowley 1991). While these so-called ‘petal displays’ are well documented in this genus, here we report for the first time the use of anthropogenic materials during such displays in two fairy-wren species.

### Study Sites

The White-winged Fairy-wren *Malurus leucopterus* observation was made on the outskirts of the City of Kalgoorlie-Boulder, Western Australia, at South Boulder waste water treatment plant (WWTP) (30°48’33”S 121°29’35”E). The area was dominated by saltbush (*Atriplex* sp.) and bluebush (*Maireana* sp.) and is interspersed with scattered *Eucalyptus* trees. There was minimal ground cover leaving the bare lateritic soils exposed between the shrubs and trees. As this was an active work site, the habitat was heavily disturbed with damaged vegetation, tyre tracks gouging deep ruts, soil heaped into mounds and ridges and there was rubbish scattered across the landscape including many pieces of white soft plastic caught on shrubs and fences.

The Red-backed Fairy-wren *M. melanocephalus* observation reported here occurred at a long-term bird ecology field site in Southeast Queensland on the western fringe of Lake Samsonvale (27°16′S, 152°51′E). The study site hosts many habitats, ranging from grassland with sparse *Eucalyptus* species to remnant old-growth rainforest and lake-edge wetlands. The dominant understory species is the invasive *Lantana camara*, which is present across habitats at our site. Three fairy-wren species are present at the site: Red-backed, Superb *M. cyaneus*, and Variegated *M. lamberti* of which most are colour-banded including approximately 200 Red-backed Fairy-wrens. Study efforts at this site began in 2012 and occur annually (except 2020–21) from July to January, and center on colour-banding and monitoring nests of all three fairy-wren species. Nests are marked using orange flagging tape tied to trees and shrubs 5–10m from nests to allow efficient navigation to nests during checks by personnel. Many broad-spanning ecological studies have been conducted at this site, mainly encompassing the colour-banded fairy-wren populations and their brood parasites (Feeney et al. 2018; Kennerley *et al*. 2019; Poje *et al*. 2019; Richardson *et al*. 2019; Hawkins *et al*. 2020; Carr *et al*. 2020; Boersma *et al*. 2023; Resendiz *et al*. 2024; Kessler *et al*. 2024; Corneliussen *et al*. 2025; Feeney et al. 2025).

### Observations

#### White-winged Fairy-wren

On 31 January 2017, JAK observed and photographed an ornamented male White-winged Fairy-wren at South Boulder WWTP carrying a piece of white material in the manner of a petal display (Figure 1). Size, structure and reflectance of the *petal-carried* object suggested it was a piece of white soft plastic, such as from a single-use carrier bag, which were numerous littering the surrounding shrubs around this work site. Additionally, the absence of white flowers in the surrounding area supported the notion that the material was a fragment of soft plastic. This individual was the only ornamented male in a group of 12 individuals, the remainder were unsexed as they were in brown-type plumage. Multiple birds in the group were heard vocalising.

**Figure 1.**
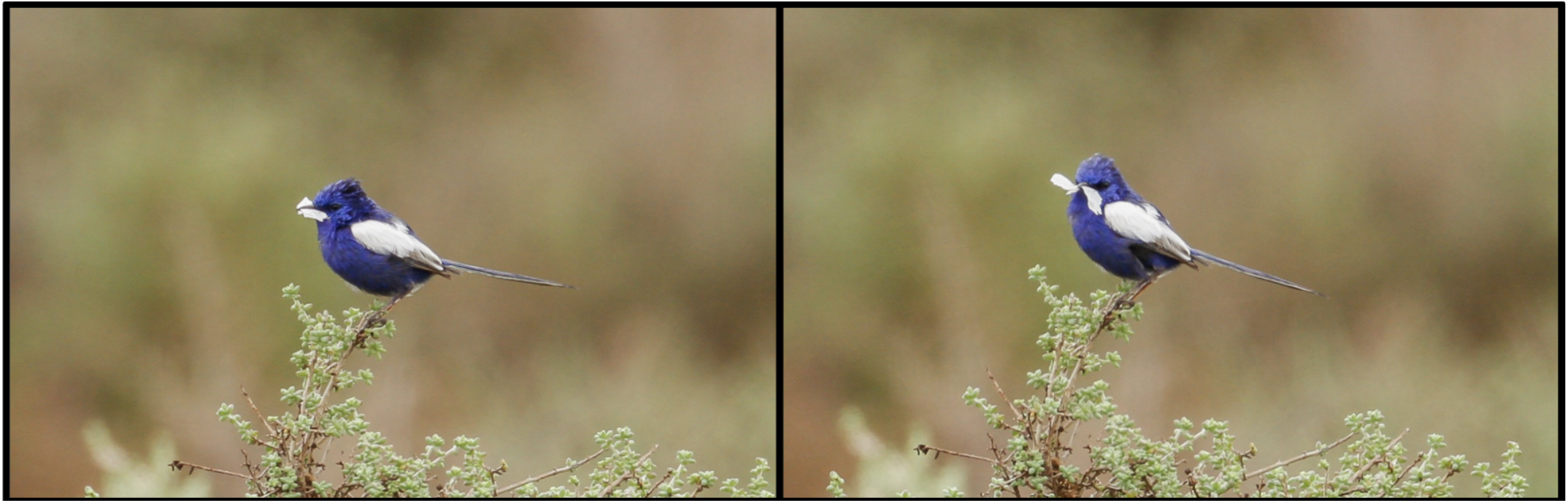
Ornamented male White-winged Fairy-wren *Malurus leucopterus* carrying a large, white, plastic object on 31 January 2017 at South Boulder waste water treatment plant, Kalgoorlie-Boulder, Western Australia (Macaulay Library catalogue numbers, left: ML115465891, right: ML115465921). *Photos by James A. Kennerley*.

#### Red-backed Fairy-wren

During a targeted mist-netting effort involving playback for Red-backed Fairy-wren on 11 August 2025 a brown-phenotype bird, likely a female, was captured in the net. Approximately 30 seconds after, JB observed an unusually long and directional flight from a male *M. melanocephalus* into the net. Upon approaching the net JB noticed an orange object lying slightly above the captured individual within the same trammel line (Figure 2) indicating that the male petal carried-carried into the net.

**Figure 2.**
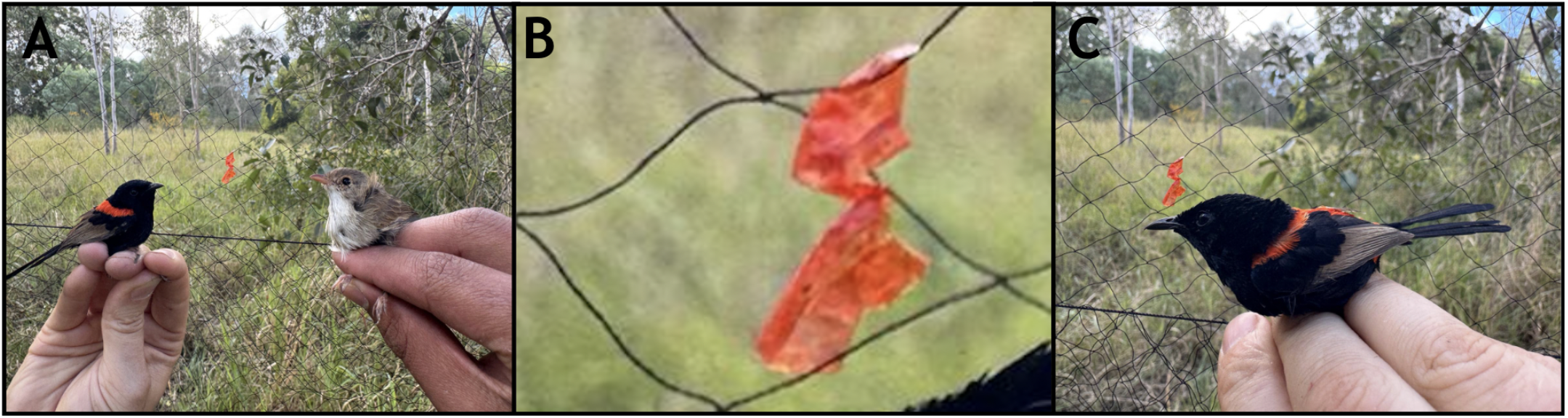
Ornamented male Red-backed Fairy-wren *Malurus melanocephalus* with brown-type Red-backed Fairy-wren alongside petal carried material (A). Enlarged photograph of the coloured flagging tape carried into the net by the ornamented male fairy-wren (B). Ornamented male Red-backed Fairy-wren held in-plane with petal-carried object (C). Photos taken on 11 August 2025 at the Samsonvale study site. Photos by *Jordan Boersma*.

JB noticed the larger size of the carried object compared to past field observations of petal-carrying behaviour. The material was stiff yet brittle, piecing apart easily after being extracted from the mistnet. Closer inspection revealed that the object was likely a piece of orange flagging tape used to mark a nest from a previous year. After processing and releasing both captured individuals, we searched to find older flagging tape for comparison. Upon examining flagging tape in the area from previous years, the colour, texture, and brittleness of orange flagging tape closely matched the object found with the individual in the net.

## Discussion

Male fairy-wrens are widely documented to exhibit petal-carrying behaviour which can include carrying flowers, dried leaves, berries, grasses, or rarely insects (Rowley 1991; Karubian and Alvarado 2003). We observed both a White-winged and Red-backed Fairy-wren apparently carrying plastic during petal displays on separate occasions. White-winged Fairy-wrens have previously been documented to carry petals in a wide range of hues from pink and yellow, to purple, blue, and white (Rowley 1991; Birdlife Australia 2023b), whereas Red-backeds are most frequently observed petal-carrying *Lantana* flower petals on the red-yellow colour spectrum (Welklin 2020; Baldassarre *et al*. 2024).

Our search for published material and media in the Macaulay Library (macaulaylibrary.org) found no previous documentation of any fairy-wren species using anthropogenic material for petal displays. Thus, to the best of our knowledge, the observations we report here of White-winged and Red-backed Fairy-wrens are the first evidence of anthropogenic material being used in a sexual display in the *Malurus* genus. Use of anthropogenic materials for an extended ornamented phenotype has been well documented in other species, such as bowerbirds. For instance, male Satin Bowerbirds *Ptilonorhynchus violaceus* commonly decorate their bowers with anthropogenically-derived blue items (Wojcieszek *et al*. 2006; Lavers *et al*. 2025). Our observations are the first records outside of bowerbirds of a bird using an artificial material for a sexual display.

Petal-carrying is known to be a primary mechanism for extra-pair copulation in Red-backed Fairy-wrens, which underlies fitness differences across male phenotypes (Baldassarre *et al*. 2016; Dowling and Webster 2017), and is likely to operate in a similar fashion in other fairy-wren species. Continued work will determine whether flagging tape petals displays increase in frequency within the three fairy-wren species at our main study site at Lake Samsonvale as this is a readily available resource at our site and offers a larger colourful material than the plant material available at our site (Appendix 1). Moreover, it is likely that other *Malurus* use anthropogenic materials, such as plastics, in petal displays given many species’ proximity to large human settlements but have been unrecognized or unreported. We hope that the ever-increasing use of participatory science platforms like eBird and iNaturalist (inaturalist.org) will allow us to better understand the frequency of these occurrences.

## Acknowledgements

We are grateful to Seqwater for access to our Lake Samsonvale study site, and to everyone who submitted photographs of petal-carrying fairy-wrens to the Macaulay Library. Jordan Karubian, Anne Peters, and Joe Welklin, who have extensive experience observing petal-carrying behaviour in Malurids helped confirm the novelty of our observations. This study was conducted with approval by Cornell IACUC (2015-0065), the Queensland Department of Environmental Science (P-PTUKI-100283615) and Seqwater (NPD20250326). We were supported by Birds Queensland, the Cornell Lab of Ornithology, Doñana Biological Station, and the Wildlife Research and Education Institute.

## APPENDIX 1

**Figure A1:**
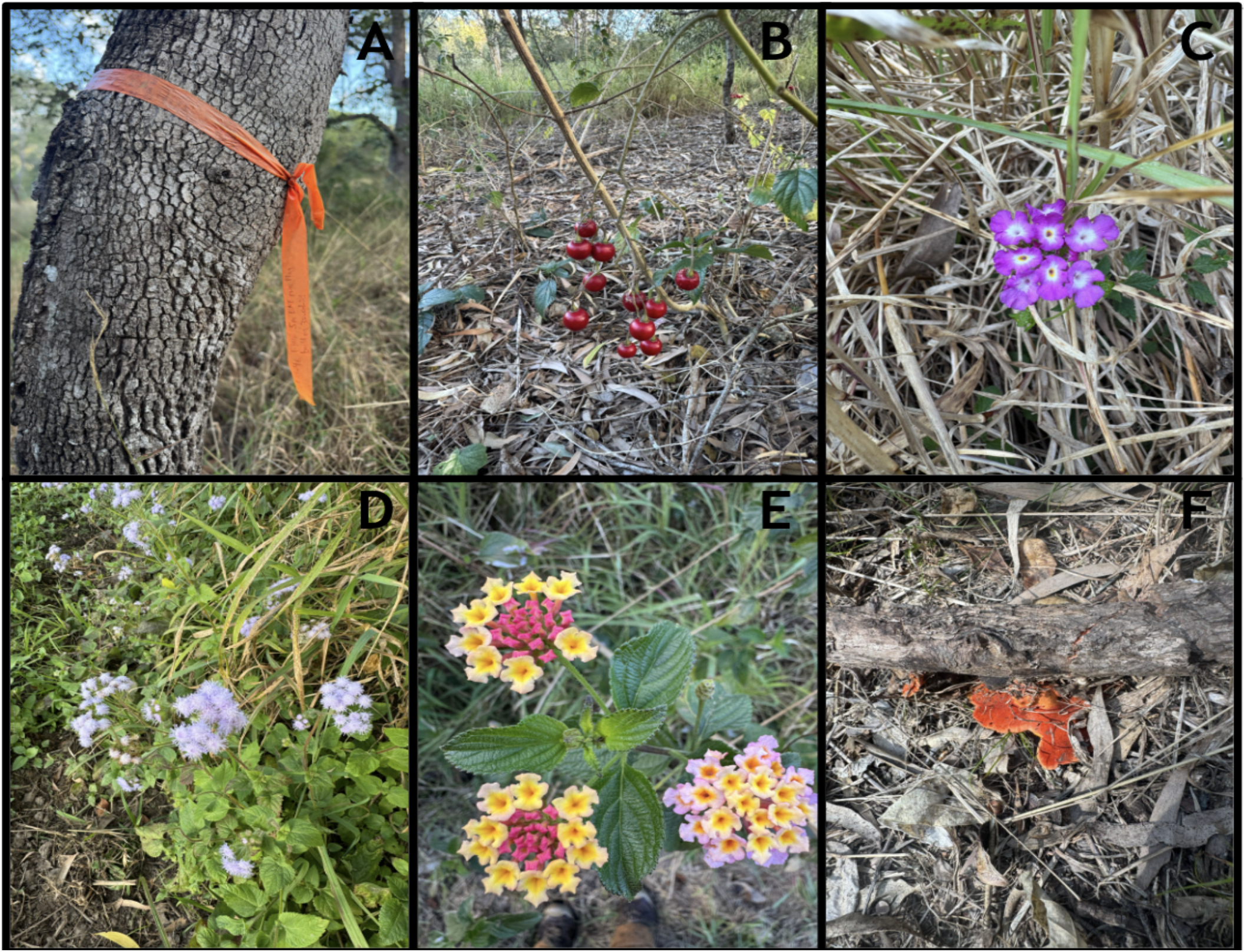
Flagging tape typically used to mark nests of *M. melanocephalus, M. lamberti, and M. cyaneus* (A). Plant and fungal matter available for petal displays at Lake Samsonvale (Panels B-F).

## References

Baldassarre DT, Greig EI, Webster MS. (2024). Red-backed Fairywren (Malurus melanocephalus), version 2.0. Birds of the World. doi:10.2173/bow.rebfai1.02

Baldassarre DT, Greig EI, Webster MS (2016). The couple that sings together stays together: duetting, aggression and extra-pair paternity in a promiscuous bird species. Biology Letters 12, 20151025. doi:10.1098/rsbl.2015.1025

Birdlife Australia (2023a). Red-backed Fairy-wren. In ‘Red-backed Fairy-wren. [Text before updates sourced from: Marchant, S. et al (eds) 1990-2006 Handbook of Australian, New Zealand and Antarctic Birds.Volume 1 to 7.] Birdlife Australia.’ (Birdlife Australia) Available at: https://hanzab.birdlife.org.au/species/red-backed-fairy-wren/ [accessed 9 September 2025]

Birdlife Australia (2023b). White-winged Fairy-wren. In ‘White-winged Fairy-wren. [Text before updates sourced from: Marchant, S. et al (eds) 1990-2006 Handbook of Australian, New Zealand and Antarctic Birds.Volume 1 to 7.] Birdlife Australia.’ (Birdlife Australia) Available at: https://hanzab.birdlife.org.au/species/white-winged-fairy-wren/ [accessed 9 September 2025]

Blamires S. (2013). Spider webs as extended phenotypes. Spiders: Morphology, Behavior and Geographic Distribution, 47–70.

Boersma J, Thrasher DJ, Welklin JF, Baldassarre DT, Feeney WE, Webster MS. (2023). Plural breeding among unrelated females and other insights on complex social structure in the cooperatively breeding Variegated Fairywren. Emu - Austral Ornithology 123, 232–243. doi:10.1080/01584197.2023.2230478

Borgia G (1985). Bower quality, number of decorations and mating success of male satin bowerbirds (Ptilonorhynchus violaceus): an experimental analysis. Animal Behaviour 33, 266–271. doi:10.1016/S0003-3472(85)80140-8

Carr HH, Kennerley JA, Richardson NM, Webster MS, Feeney WE (2020). First record of black feathering in a female Red-backed Fairy-wren Malurus melanocephalus under natural conditions. Australian Field Ornithology 37, 150–154.

Corneliussen L, Kennerley JA, Zamora M, Lucille S, Jack R, Monteith M, Partridge S, Webster MS, Boersma J, Feeney WE (2025). Repeated nest failure precedes group dissolution and joining a neighbouring group by a female Variegated Fairy-wren Malurus lamberti. Australian Field Ornithology 42, 00–00. doi:10.20938/afo42000000

Dawkins R (1982). ‘The extended phenotype’. (Oxford university press Oxford)

Dowling J, Webster MS (2017). Working with what you’ve got: unattractive males show greater mate-guarding effort in a duetting songbird. Biology Letters 13, 20160682. doi:10.1098/rsbl.2016.0682

Dunn, P. O., & Cockburn, A (1999). Extrapair mate choice and honest signaling in cooperatively breeding superb fairy-wrens. Evolution, 53(3), 938–946.

Feeney WE, Ryan TA, Kennerley J, Poje C, Clarke L, Scheuering M, Webster MS (2018) A photographic guide for ageing nestlings of two species of Australian brood parasitic cuckoo: the fan-tailed (Cacomantis flabelliformis) and Horsfield’s bronze (Chalcites basalis) cuckoos.Australian Field Ornithology 35: 8-12. doi/abs/10.3316/informit.400247613632857

Feeney WE, Kennerley JA, Wheatcroft D, Liang W, Lamb JB, Teunissen N, Lawson SL, Enos JK, Zhou B, Poje C, Richardson NM, Ryan TA, Zamora M, Cowan Z-L, Brooker RM, Lamb JB, Attwood M, Gloag R, Fiorini VD, Gill SA, Peters A, Honza M, Spottiswoode CN, Hauber ME, Manica A, Webster MS, Blasi D (In Press) Learned use of an ancient sound-meaning association in birds. Nature Ecology & Evolution

Hawkins CE, Ritrovato IT, Swaddle JP (2020). Traffic noise alters individual social connectivity, but not space-use, of Red-backed Fairywrens. Emu - Austral Ornithology 120, 313–321. doi:10.1080/01584197.2020.1830706

Karubian J, Alvarado A (2003). Testing the function of petal-carrying in the Red-backed Fairy-wren (Malurus melanocephalus). Emu - Austral Ornithology 103, 87–92. doi:10.1071/MU01063

Kennerley JA, Grundler MR, Richardson NM, Marsh M, Grayum J, Feeney WE (2019). Observations on the behaviour and ecology of the Pallid Cuckoo Heteroscenes pallidus in south-eastern Queensland. Australian Field Ornithology 36, 109–115. doi:10.20938/afo36109115

Kessler WP, Kennerley JA, Resendiz E, Ditzel PC, Buckley ER, Huff CE, Poje C, Coleman JT, Boersma J, Webster MS, Feeney WE (2024). Use of a communal display area by Rufous Whistlers Pachycephala rufiventris. Australian Field Ornithology 41, 204–205. doi:10.20938/afo41204205

Lavers JL, Fidler AL, Charlton-Howard H (2025). Anthropogenic pollution is widespread in Great Bowerbird bowers in northern Australia. Microplastics and Nanoplastics 5, 27. doi:10.1186/s43591-025-00133-w

Moller AP, Pomiankowski A (1993). Why have birds got multiple sexual ornaments? Behavioral Ecology and Sociobiology 32, 167–176. doi:10.1007/BF00173774

Moreno J, Soler M, Møller AP, Linden M (1994). The function of stone carrying in the black wheatear, Oenanthe leucura. Animal Behaviour 47, 1297–1309. doi:10.1006/anbe.1994.1178

Poje C, Kennerley JA, Richardson NM, Cowan Z-L, Grundler MR, Marsh M, Feeney WE (2019). Notes on the parasitic ecology of newly-fledged Fan-tailed Cuckoos Cacomantis flabelliformis and their White-browed Scrubwren Sericornis frontalis hosts in south-east Queensland. The Sunbird 48, 162–166.

Resendiz E, Ditzel PC, Kessler WP, Buckley ER, Huff CE, Poje C, Lamb JB, Kennerley JA, Coleman JT, Boersma J, Webster MS, Feeney WE (2024). A photographic guide for determining egg incubation stage in the Superb Fairy-wren Malurus cyaneus. Australian Field Ornithology 41, 150–154. doi:10.20938/afo41150154

Richardson NM, Kennerley JA, Feeney WE (2019). First record of intraspecific adoption by a female Superb Fairy-wren, Malurus cyaneus. The Sunbird 48, 159–161.

Rowley I (1991). Petal-carrying by fairy-wrens of the genus Malurus. Australian Field Ornithology 14. Available at: https://afo.birdlife.org.au/afo/index.php/afo/article/view/844 [accessed 9 September 2025]

Schaedelin FC, Taborsky M (2009). Extended phenotypes as signals. Biological Reviews 84, 293–313. doi:10.1111/j.1469-185X.2008.00075.x

Turner JS (2004). Extended Phenotypes and Extended Organisms. Biology and Philosophy 19, 327–352. doi:10.1023/B:BIPH.0000036115.65522.a1

Webster, M. S., Tarvin, K. A., Tuttle, E. M., & Pruett-Jones, S (2007). Promiscuity drives sexual selection in a socially monogamous bird. Evolution, 61(9), 2205–2211.

Webster MS, Varian CW, Karubian J (2008). Plumage color and reproduction in the red-backed fairy-wren: Why be a dull breeder? Behavioral Ecology 19, 517–524. doi:10.1093/beheco/arn015

Welklin JF (2020). Social and environmental causes and reproductive consequences of ornamented plumage in the Red-backed Fairywren (Malurus melanocephalus). Available at: https://hdl.handle.net/1813/103273 [accessed 9 September 2025]

Wells DA (2015). The extended phenotype(s): a comparison with niche construction theory. Biology & Philosophy 30, 547–567. doi:10.1007/s10539-015-9476-0

Wojcieszek JM, Nicholls JA, Marshall NJ, Goldizen AW (2006). Theft of bower decorations among male Satin Bowerbirds (Ptilonorhynchus violaceus): why are some decorations more popular than others? Emu - Austral Ornithology 106, 175–180. doi:10.1071/MU05047

